# Mining unknown porcine protein isoforms by tissue-based map of proteome enhances the pig genome annotation

**DOI:** 10.1101/391466

**Authors:** Pengju Zhao, Xianrui Zheng, Ying Yu, Zhuocheng Hou, Chenguang Diao, Haifei Wang, Huimin Kang, Chao Ning, Junhui Li, Wen Feng, Wen Wang, George E. Liu, Bugao Li, Jacqueline Smith, Yangzom Chamba, Jian-Feng Liu

**Author notes:** Corresponding author: Jian-Feng Liu, Ph.D., China Agricultural University (West Campus) College of Animal Science and Technology Room 455, 2 Yuanmingyuan West Road, Beijing 100193, China. These authors contributed equally to this work (Pengju Zhao, Xianrui Zheng).

## Abstract

A lack of the complete pig proteome has left a gap in our knowledge of the pig genome and has restricted the feasibility of using pigs as a biomedical model. We developed the tissue-based proteome maps using 34 major normal pig tissues. A total of 7,319 unknown protein isoforms were identified and systematically characterized, including 3,703 novel protein isoforms, 669 protein isoforms from 460 genes symbolized beginning with LOC, and 2,947 protein isoforms without clear NCBI annotation in current pig reference genome. These newly identified protein isoforms were functionally annotated through profiling the pig transcriptome with high-throughput RNA sequencing (RNA-seq) of the same pig tissues, further improving the genome annotation of corresponding protein coding genes. Combining the well-annotated genes that having parallel expression pattern and subcellular witness, we predicted the tissue related subcellular components and potential function for these unknown proteins. Finally, we mined 3,656 orthologous genes for 49.95% of unknown protein isoforms across multiple species, referring to 65 KEGG pathways and 25 disease signaling pathways. These findings provided valuable insights and a rich resource for enhancing studies of pig genomics and biology as well as biomedical model application to human medicine.

## Background

The domestic pig (Sus *scrofa*) is one of the most popular livestock species predominately raised for human consumption worldwide. Besides its socio-economic importance, pig has been generally recognized as a valuable model species for studying human biology and disease due to its striking resemblances with humans in anatomy, physiology and genome sequence(Ekser et al. 2015; Cooper et al. 2016). To date, many porcine relevant biomedical models have been created for exploring etiology, pathogenesis and treatment of a wide range of human diseases, *e.g.*, Parkinson disease(Bjarkam et al. 2008), obesity(Pedersen et al. 2013), brain disorder(Lind et al. 2007), cardiovascular, atherosclerotic disease(Agarwala et al. 2013) and Huntington’s disease(Yan et al. 2018), *etc*. Furthermore, pigs and humans share similarities in the size of their organs, making pig organs potential candidates for porcine-to-human xenotransplantation(Cooper 2012; Li et al. 2016). Recently major efforts have been devoted to the development of tools for further enhancing the value of pigs as a biomedical model for human medicine as well as its role in meat production. Of essential significance is the completion of the assembly of the pig genome sequence (*Sus scrofa11.1*) in recent time. It provides researchers with a vast amount of genomic information, facilitating characterization of individual pig genome as well as genome comparison between pigs and humans.

With the progress of large-scale genome projects, such as ENCODE(Consortium 2012) and Human Proteome Projects(Legrain et al. 2011), many genes have been annotated at the RNA and protein levels, and diverse regulatory elements across the human genome were systematically characterized. This creates great opportunities for exploring how genetic variation underlies complex human phenotypes(Maher 2012). In particular, a spate of groundbreaking studies were succeeded in building high-resolution maps of the proteome(Kim et al. 2014; Wilhelm et al. 2014; Uhlen et al. 2015) in a variety of human tissues and cells. Findings from these studies greatly facilitate the functional annotation of the genome at multiple-omic levels and further improve the understanding of complexity of human phenotypes.

Compared with humans, however, studies of pig proteome are very limited(Chen et al. 2015; Fischer et al. 2015). In particular, in-depth identification and characterization of the proteome maps of the pig genome across a broad variety of pig tissues is not yet available. To date, the leading protein database UniProtKB comprised around 1,419 reviewed and 34,201 unreviewed pig proteins in Swiss-Prot and TrEMBL respectively. It is far less than the numbers of entries in Swiss-Prot (20,215 proteins) and TrEMBL (159,615 proteins) corresponding to human proteome data. Although the recent update of the pig PeptideAtlas presented 7,139 protein canonical identifications from 25 tissues and three body fluid(Hesselager et al. 2016), this is still a limited promotion to whole pig proteome research. In fact, a large number of unreviewed and PeptideAtlas-identified pig proteins were not well annotated in current genome (*Sus scrofa11.1*) due to lack of specific genomic locations and the corresponding assembled RNA transcripts. This suggests that there are still plenty of poorly annotated proteins that are not identified and characterized in previous pig studies. Besides, even if the annotated pig protein-coding genes (PCGs), nearly 20% of which were symbolized beginning with LOC — the orthologs and function of genes have not been determined — that also appeared one of the key limitations of pig gene set enrichment analysis. The absence of completive maps for the pig proteome triggers a substantial bottleneck in the progress of refining pig genome annotation and even hinders systematic comparison of omics data between humans and pigs.

Therefore, considering the potential contribution to developing pig proteomic atlases, we conducted in-depth characterization of pig proteome across 34 histologically normal tissues using high-resolution mass spectrometry. Accordingly, we exploited the novel protein firstly identified herein, poorly annotated proteins, and LOC proteins and defined these as the pig unknown proteins. These unknown proteins were mapped to the latest pig genome (*Sus scrofa11.1*) for confirming their available genomic locations. We then constructed pig transcriptomic atlas and subcellular characterization for these unknown protein isoforms to infer their connections with the specific function of tissues. Finally, systematically comparing the orthologous relationship of these unknown proteins with other multiple species, we further predicted the potential function of these unknown protein isoforms to ensure their availability in future relevant studies. Findings herein will benefit studies and development of pig genome and will allow further investigation of swine gene function and networks of particular interest to the scientific community.

## Results

### Tissue-based map of the pig proteome

We explored proteome from 34 pig tissue (Figure 1a) samples using liquid chromatography tandem mass spectrometry (LC-MS/MS). We performed *in silico* analyses (Figure 1b) to construct the whole landscape of the pig proteome with a view to furthering pig biological research and human medical studies. The resulting proteome data involved a total number of 21,681,643 MS/MS spectra produced from 680 LC-MS/MS runs (20 runs per tissue).

**Figure 1.**
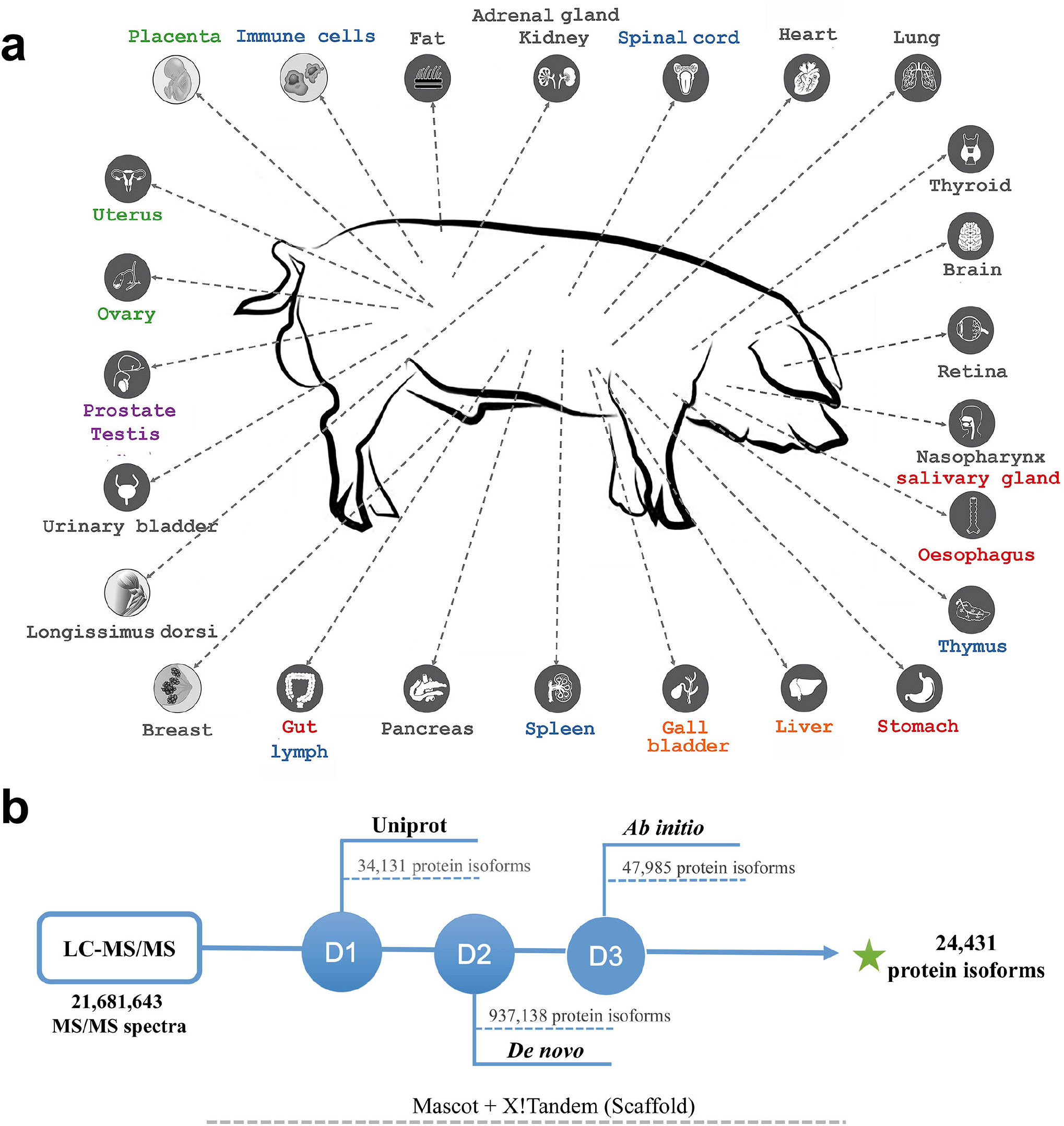
Overview of pig transcriptome-based annotation. **a.** 34 pig tissues analyzed in this study. 34 representative normal pig tissues were selected as the resource of proteome and transcriptome for exploring convincing evidence of putative PCGs, where A and I respectively represent adult and infancy pig tissue. **b.** The custom pipeline for proteome-based annotation. Four protein database were used for protein searching based on Mascot and X!Tandem software with the same criteria.

To exploit convincing peptide evidence for all putative PCGs in the pig genome, we searched the raw MS/MS data by Mascot(Perkins et al. 1999) against multiple protein databases. These included the primary pig database of UniProt(UniProt 2015) for the initial search and two custom-developed databases for sequential searches of unmatched spectra, *i.e.*, (1) RNA-seq-based *de novo* assembly transcriptomic database which included the RNA-seq data generated from the 34 tissues in this study, 1.08 Gb data from an external public expressed sequence tag (EST) database and 953.57Gb from publicly available RNA-seq data (Materials and Methods), (2) a six-frame-translated pig genome database. Those corresponding matched spectra extracted from each subset of databases were re-searched against the same database by X!Tandem(Craig and Beavis 2003) for further filtration to produce the final 5,082,599 peptide spectrum matches (PSMs). Subsequently, Scaffold (version Scaffold_4.4.5, Proteome Software Inc., Portland, OR) was run for MS/MS-based peptide and protein identification, both using the global false discovery rate (FDR) criterion of 0.01.

Totally, we identified 212,154 non-redundant peptides with a median number of 8 unique peptides per gene (Quality assessment of protein identification is shown in Figure S1-14). Comparison of identified peptides with the largest pig peptide resource PeptideAtlas (http://www.peptideatlas.org/) showed that 49,144 out of 87,909 curated peptides (56%) were confirmed by our identification. The peptides we detected greatly outnumbered those deposited in PeptideAtlas, with a major fraction (77%) found to be novel. A total of 24,431 protein isoforms with median sequence coverage of 30.32% were determined by Scaffold, which corresponded to 19,914 PCGs. To ascertain whether our protein identifications included a reasonable false positive error rate, we additionally validated 31 proteins from different proteogenomic categories. By comparing MS/MS spectra from 71 synthetic peptides with those obtained from our analysis of pig tissues, we obtained 100% validation (Table S1; Supplemental file 1).

### Identification and characterization of unknown pig proteins

Classifying all of 24,431 identified protein isoforms (Figure 2a), we found 16,738 (68.51%) protein isoforms were confirmed by the Uniprot protein evidence, 9,204 (37.67%) protein isoforms had evidence from pig PeptideAtlas(Hesselager et al. 2016), 17,781 (72.78%) protein isoforms were included in NCBI protein database, and 7,910 (32.38%) were supported by all of them. Of all confirmed protein isoforms, 17,781 (85.78%) protein isoforms according to 11,308 PCGs were included in known NCBI annotation, 669 protein isoforms according to 460 PCGs were annotated in the pig genome but classified as uncharacterized LOC genes, and 2,947 protein isoforms remain a lack of NCBI annotation support in the pig genome (Figure 2b). The rest of 3,703 protein isoforms were identified by MS/MS data for the first time in this study, which can be considered as potential novel proteins.

**Figure 2.**
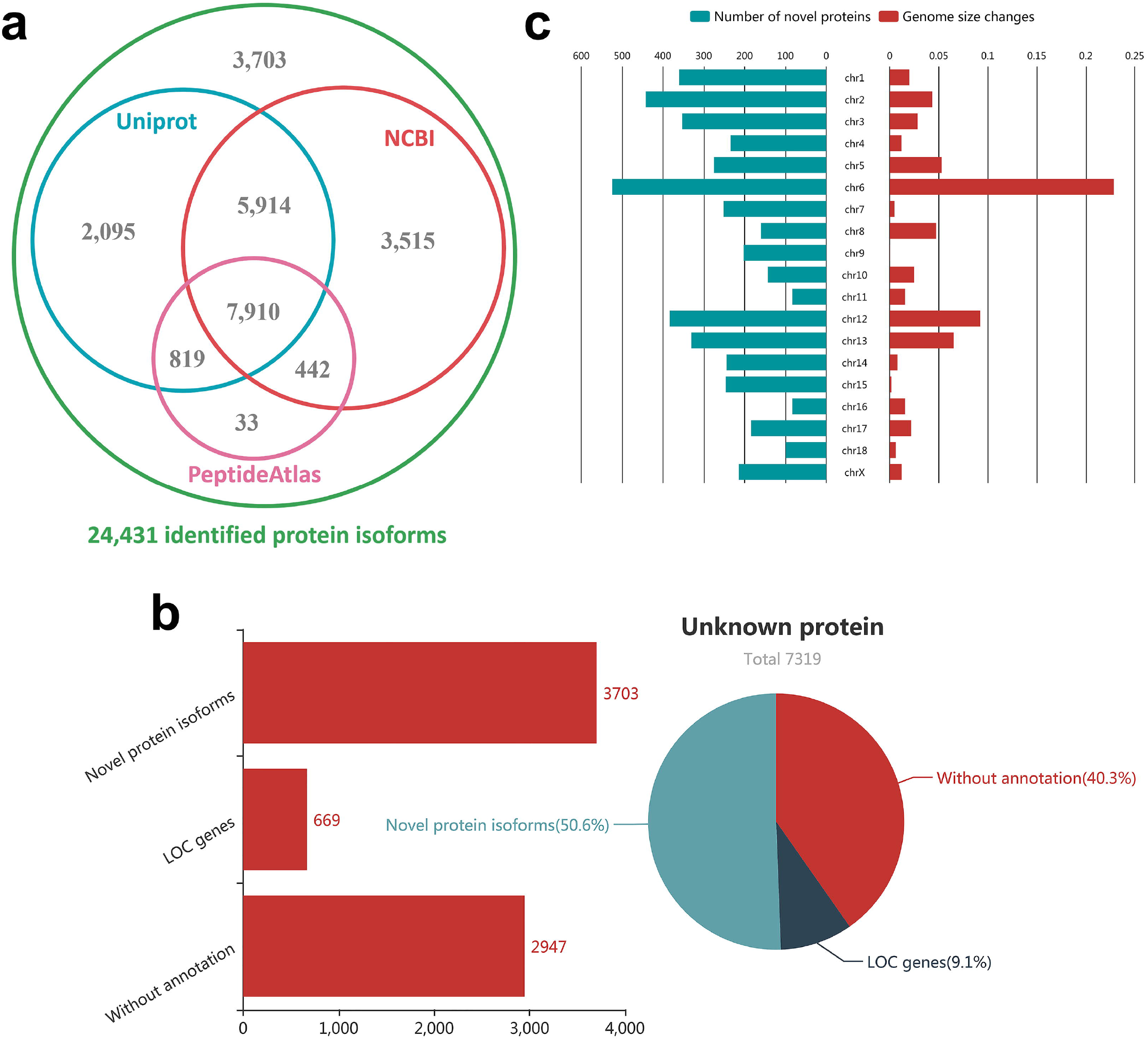
Characterization of unknown pig protein isoforms. **a.** Confirmation of 24,431 identified protein isoforms by other pig protein databases. **b.** Classification of unknown pig protein isoforms. Bar chart and pie chart respectively show the numbers and percents of three categories in 7,319 unknown pig protein isoforms. **c.** Relationship between the improvement of genome quality and novel proteins.

To further enhance the annotation of PCGs for the current pig genome, we systematically characterize these 7,319 feature or/and location unknown protein isoforms detected in current study (*i.e.*, 669 protein isoforms of LOC genes, 2,947 protein isoforms without genomic location annotation and 3,703 protein novel isoforms firstly identified herein). Considering only 9.14 % of protein isoforms (LOC genes) had the available genomic locations, we mapped the rest of 6,650 unknown protein isoforms to the pig reference genome (*Sus scrofa11.1*) by MAKER annotation workflow (Cantarel et al. 2008). First, the low-complexity repeats of pig reference genome were soft-masked by RepeatMasker. Totally 6,650 out of 7,319 unknown protein isoforms (non-LOC genes) were aligned to the masked reference genome by BLAST(Mount 2007). Sequentially, Exonerate(Slater and Birney 2005) was run to realign and polish the exon–intron boundaries of the unknown gene with the splice-site aware alignment algorithm. A total of 5,027 (75.6%) unknown protein isoforms (non-LOC genes) were successfully aligned to the reference genome with the sequence identity > 95% and similarity > 95% (2,381 with the 100% identity and 100% similarity), including 4,819 assigned to chromosomes and 208 resided on 33 unplaced scaffolds. More interestingly, we found that the proportion of novel proteins mapped in respective chromosomes was related to the levels of genomic improvement from *Sus scrofa10.2* to *Sus scrofa11.1* for different chromosomes (Figure 2c, R^2^ = 0.67, *P* = 0.0015). This demonstrated that these unknown proteins, especially the novel proteins, were actually ignored in current pig genome annotation since most of previous studies have been limited to *Sus scrofa10.2* genome and fewer tissues.

Comparison of these unknown proteins isoforms with the well-annotated proteins revealed that, a major fraction of unknown protein isoforms (40.29%), especially the novel protein isoforms (53.07%), were merely identified in a single tissue that far more than well-annotated protein isoforms. It can be speculated that most of novel protein isoforms were more likely tissue specificity, resulting in the neglect of proteins in previous studies. Additionally, further analysis of the reliability for these unknown proteins, we found a major fraction of them (52.55%) were regarded as the abundant proteins that have more than ten spectral counts(Zhou et al. 2012). Particularly, although these novel protein isoforms were first identified in this study, almost 65.75% of all were supported by a high spectral count of > 5, indicating that the identification of these novel protein isoforms significantly enhances the current pig protein database with convincing evidence.

### Expression landscape of unknown protein isoforms by profiling pig transcriptome

To further probe potential function of unknown protein isoforms, we characterized the expression landscape of unknown protein isoforms by high-throughput RNA sequencing (RNA-seq) of the 34 identical tissue samples as in the LC-MS/MS analyses. Compared to the label-free LC-MS/MS method that mainly applied to protein identification, RNA-seq was able to better reflect the gene expression level in the organism.

Approximately 1,495 million paired-end reads (376.7G bases per tissue) were obtained through sequencing 116 strand-specific paired-end RNA libraries, of which 1,230 million were mapped to the pig genome (*Sus scrofa* 11.1) with an overall pair alignment rate of 88.29% (Table S2). As expected, a total number of 2,486,239 transcripts (the FPKM > 0.1 in at least one tissue) corresponding to 29,270 genes were then assembled and quantified across all tissues, which contained 5,250 annotated transcripts corresponding to 3,486 known noncoding genes, 7,595 potentially novel alternatively spliced transcripts corresponding to 2,421 known noncoding genes, 55,328 annotated transcripts corresponding to 20,401 PCGs, 136,537 potentially novel alternatively spliced transcripts corresponding to 15,385 PCGs, and 2,281,529 newly assembled transcripts corresponding to 26,493 genes in the pig genome without annotation information. These findings clearly increased the average number of isoforms per gene (Human-NCBI: 7.27, Pig-NCBI: 2.75, Pig-Identified: 6.60) compared with existing gene annotation in NCBI (Figure 3a).

**Figure 3.**
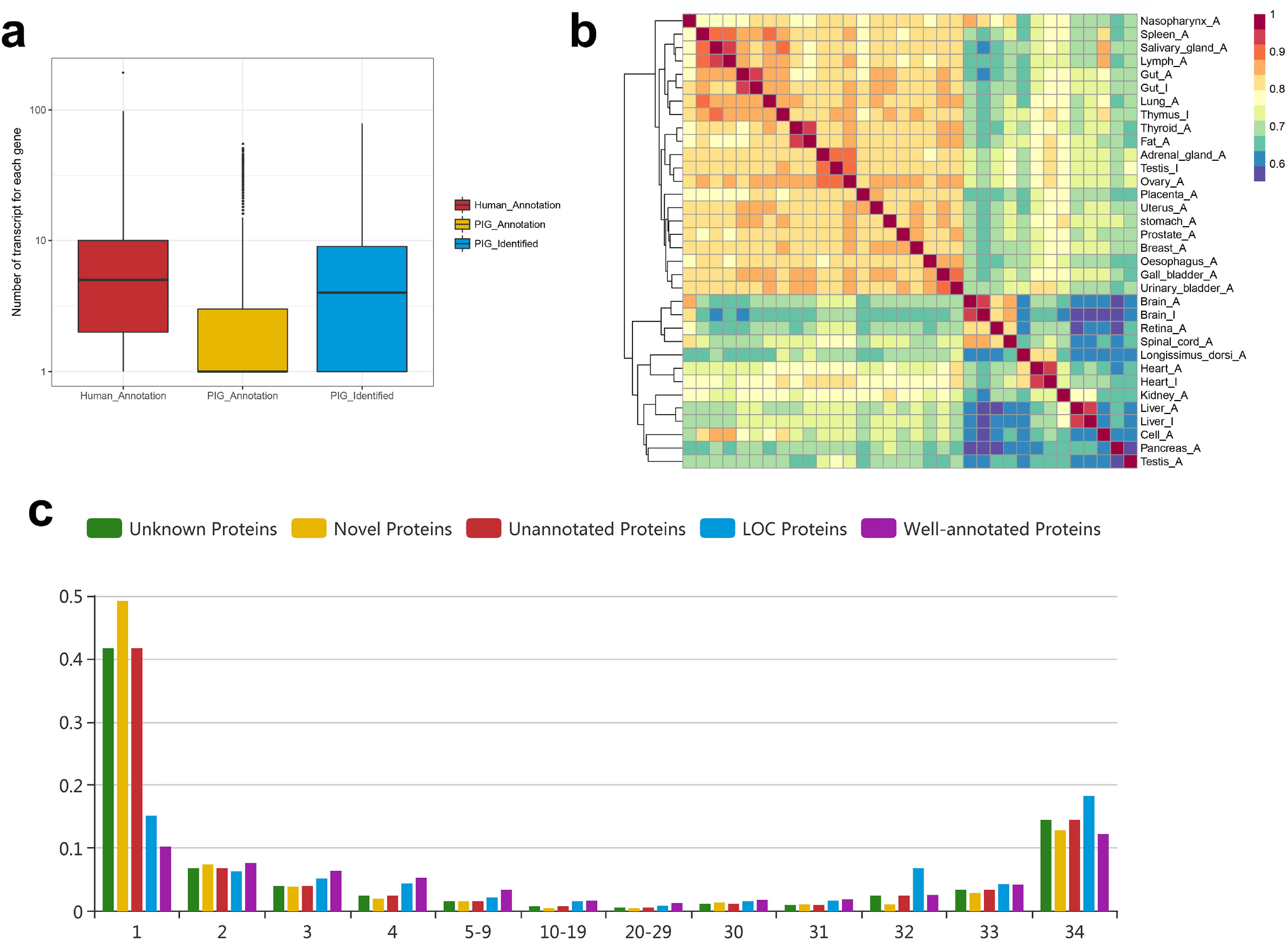
The pig transcriptome in unknown protein isoforms. **a.** Comparison of number of isoforms expressed per gene between humans and pigs. The box plots compare the number of isoforms expressed per gene within three transcript sets, including known human Ensembl set, known pig Ensembl set and the newly identified set of this study. **b.** The heatmap for Pearson correlation between 34 tissues. The heatmap was used to reveal the pairwise correlation between all 34 pig tissues. **c.** Bar chart for tissue-based transcriptomic evidence of unknown protein isoforms. The x-axis represents the numbers of tissue and the y-axis represents the numbers of protein.

On the basis of all the currently well-annotated genes, we constructed a tissue similarity map using hierarchical clustering based on the Pearson correlation across the 34 tissues. As shown in Figure 3b, (with the exception of three obvious outliers - adult testis, pancreas and peripheral blood mononuclear cells (PBMC)), data clustered into multiple known connected groups: liver and kidney, muscular system (longissimus dorsi and heart), nervous system (retina, brain and spinal cord), adult immune organs (spleen, salivary gland and lymph), bladder tissue (urinary bladder, gall bladder and oesophagus). These results showed the expected biology that had a similar expression profile to that of human tissues(Uhlen et al. 2015), reflecting the biological similarity between human and pig, as well as the reliability of transcripts we constructed.

Intriguingly, we observed a total of 47.72% (3,493) of unknown protein isoforms were successfully confirmed by the transcripts constructed herein, which offered a detailed view of the understanding of unknown proteins. Considering the comparison of unknown protein isoforms with the potential low-expression levels in different tissue, we applied zFPKM normalization method (Hart et al. 2013) to generate high-confidence estimates of gene expression. The observed zFPKM range of unknown protein isoforms expression ranged from −3 to 19.89, having on lower average expression levels (zFPKM=2.53), especially the novel protein isoforms (zFPKM=2.20), than well-annotated PCGs (zFPKM=3.62). Besides, we also found that these unknown protein isoforms (average 11.7 tissues) were tend to be expressed in less tissues than well-annotated PCGs (average 21.3 tissues), and nearly 41.6% (*n*=1453) of unknown protein isoforms were only identified in single tissues. The results showed that the previously incomplete annotation of these unknown protein isoforms were more likely due to their specific expression characteristics (Figure 3c).

Screening the protein isoforms expression patterns in each tissue, we found that the majority expression of transcripts were dominated by the expression of a small proportion of genes in all of the investigated tissues (Table S3). Specifically, the adult pig tissues of prostate, longissimus dorsi, pancreas, gall bladder, *etc.*, had the least complex transcriptome, with 50% expression of the transcripts coming from a few highly expressed genes (3 to 8 transcripts). In contrast, the reproductive tissues (uterus, testis and ovary), expressed more complex transcriptome, with a large number of genes expressed. Similar patterns have also been reported in human tissue transcriptome studies(Mele et al. 2015). It was surprising that 203 unknown protein isoforms were potentially associated with 148 (13.98%) highly expressed genes, suggesting these unknown protein isoforms play an important role in basic function among tissues or organs.

### Prediction of unknown proteins function from pig transcriptome

Several approaches for systematic analysis of gene expression across different tissues have found that gene expression patterns were usually associated with their biological functions, as well as genes with the similar functions are more likely to exhibit similar expression patterns (Zheng-Bradley et al. 2010). Implementing the similar classification criteria for human genes(Uhlen et al. 2015) into the RNA-seq data generated from the multiple pig tissues herein, we classified all 23,887 putative NCBI genes (18,377 PCGs) corresponding to well-annotated 60,578 transcripts and 3,493 unknown protein isoforms into three categories for exhibiting their expression features. The number of tissue-enriched genes, group-enriched genes and ubiquitously expressed genes are also displayed as a network plot to show an overview of pig PCGs (Figure 4a).

**Figure 4.**
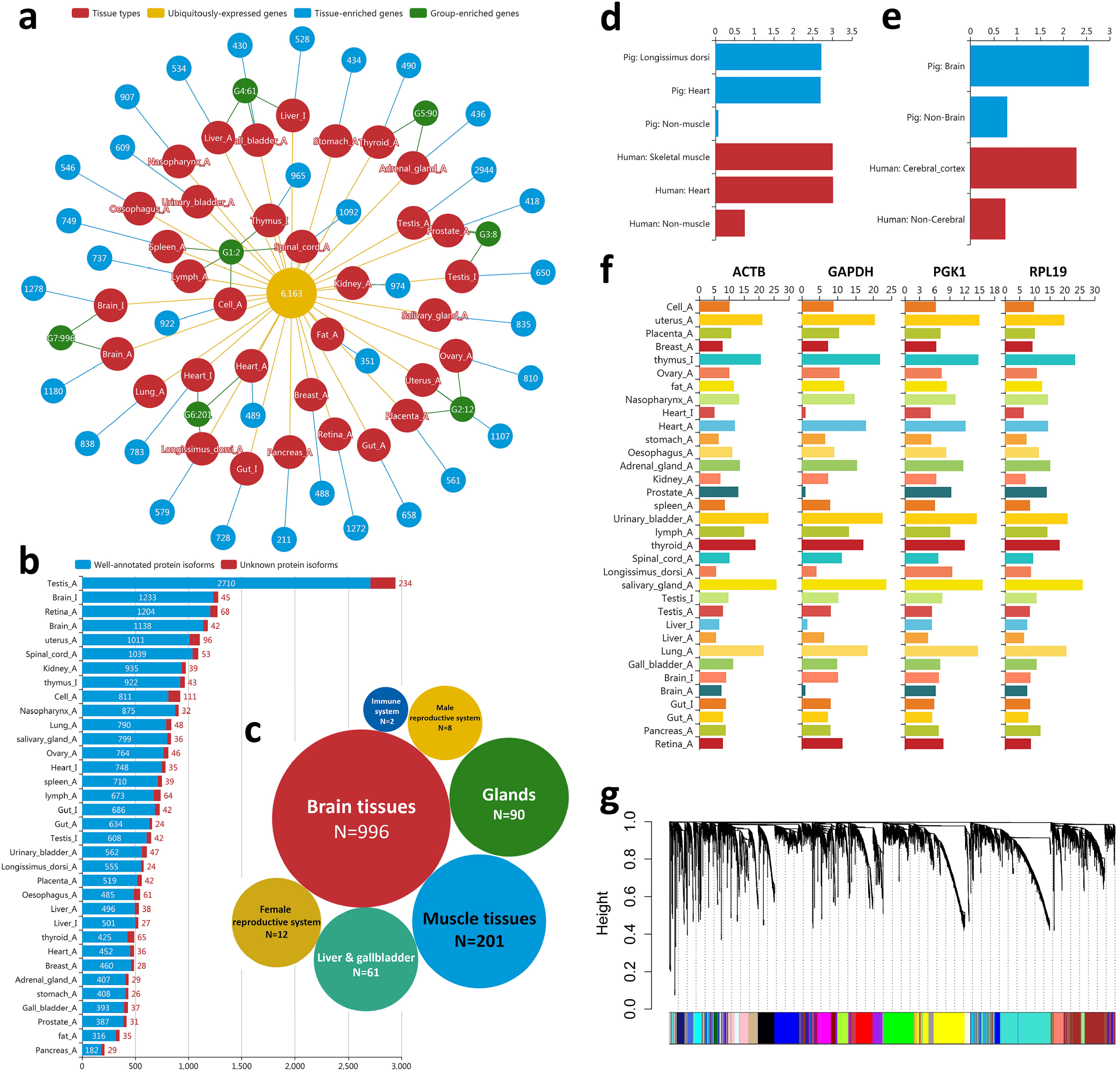
Expression landscape in pig transcriptome. **a.** The network plot for the overview of pig PCGs. The red nodes represent the types of tissue. Yellow, blue and green nodes respectively represent the number of the gene that expressed in all tissues, tissue-enriched and group-enriched. Where, G1-47 respectively means immune organs, female reproductive system, male reproductive system, Liver and gall bladder, Adrenal gland and thymus gland, Muscle tissues, Brain tissues. **b.** Numbers of tissue-enriched isoforms for known and unknown protein isoforms. **c.** Numbers of group-enriched isoforms in different tissue groups. **d.** The group-enriched expression of *MYL3* gene in muscular system. The gene levels (FPKM) for *MYL3* gene from different tissue categories (muscular system and non-muscular system) between humans and pigs. **e.** The group-enriched expression of *ENC1* gene in brain tissue. The gene levels (FPKM) for *ENC1* gene from different tissue categories (brain and brain tissue) between humans and pigs. **f.** Expression landscape of ubiquitously expressed genes in 34 tissues. **g.** Hierarchical cluster tree for all ubiquitously expressed genes. 24 modules correspond to branches are labelled by 24 different colours.

In multicellular organisms, genes expressed in a few tissue types are thought to be tissue-enriched genes which have tissue-specific related functions. We observed 8,482 (14%) well-annotated transcripts (5,592 genes) and 1,453 (41.6%) unknown protein isoforms that have a specific expression in a particular tissue, as well as 16,356 (27%) well-annotated transcripts (9,726 genes) and 241 (6.9%) unknown protein isoforms being expressed at least 5-fold higher at the zFPKM level in one tissue compared to the tissue with a second highest expression. Similarly to previous studies in humans(Uhlen et al. 2015) (Figure 4b), the largest number of tissues enriched genes were detected in the adult testis, followed by infancy brain, retina and adult brain. The results reflected that the tissues with complex biological processes usually have more tissue-enriched genes, and these tissue-enriched genes were strongly associated with the function of the corresponding tissues. This can be is exemplified by the *RHO* (Rhodopsin) gene that was enriched in retina and was proven to play important roles in retinal pigments(Yu et al. 2016). This demonstrates that the tissue specificity can not only confirm the biological characteristics of known genes but also predict basic function of undefined genes in pigs. Accordingly, we successfully updated 1,694 tissue-enriched unknown protein isoforms to further explain the functional differences among tissues.

Apart from the gene observed in tissue specificity, some group-enriched genes over-represented in the group of tissues/organs that work together to perform closely related functions. Accordingly, we found a total of 1,318 (2.18%) well-annotated transcripts (948 genes) and 52 (1.49%) unknown protein isoforms were detected and could be grouped into seven types of tissue (Figure 4c). The largest fraction (72.7%) of group-enriched genes belonged to the brain tissues (14.7%), followed by the muscular system (cardiac muscle, longissimus dorsi), adrenal and thymus gland (6.6 %), liver and gallbladder (4.5 %). Generally, these group-enriched genes have potential role in biological system function, and this expression patterns were usually shown between different species. As exemplified by the group-enriched expression of *MYL3* (myosin light chain 3, a known myosin component) (Figure 4d) and *ENC1* (Ectodermal-Neural Cortex 1, involved in mediating uptake of synaptic material) (Figure 4e). Both of these genes indicated a similar expression in the muscular system and in brain tissue between humans and pigs. Therefore, 52 of unknown protein isoforms will be the valuable resources that expected to further enrich the functional and comparative genomics between pig and human.

Specifically, we found 5,656 well-annotated transcripts corresponding to 5,147 (21.55%) NCBI genes expressed in all pig tissues. Among these genes, a variety of known “housekeeping” genes such as *ACTB, GAPDH, PGK1, RPL19, etc*. (Figure 4f) are usually intracellular and tend to be functionally essential to cell subsistence that involved in metabolism, transcription, and RNA processing or translation (Ramskold et al. 2009). Interestingly, 507 (14.5%) of unknown protein isoforms were detected as the ubiquitously expressed genes, suggesting that the findings of these unknown protein isoforms offered the important supplement to pig genomic annotation. To characterize the set of ubiquitously expression of these unknown genes identified herein, we constructed a co-expression network heatmap that consisted of 24 blocks for assessing ubiquitously expressed gene co-expression interactions across all pig tissues (Figure 4g). Obviously, these unknown protein isoforms have potentially functional connections with the well-annotated gene that in the same blocks (Table S4), which can be explained by those genes within modules of a co-expression network may be involved in similar or related pathways or biological processes (Liang et al. 2014).

### Subcellular characterization of the unknown pig proteome

Proteins with different subcellular locations usually play different roles in physiological and pathological processes. To characterize these unknown pig proteins at the subcellular level, we performed a proteome-wide subcellular classification for all identified pig protein isoforms (*n* = 24,431) based on the existing prediction methods(Uhlen et al. 2015) (as described in Materials and Methods). We found a major fraction (72.66%) of pig protein isoforms were predicted to be soluble protein isoforms, followed by 21.55% of membrane protein isoforms and 5.79% of secreted protein isoforms (Figure 5a; Table S5). For an in-depth comparative analysis on PCGs, we further clustered all available protein isoforms into four base categories including 14,890 soluble proteins, 3,924 membrane proteins, 1,053 secreted proteins, as well as 47 membrane & secreted proteins (Table S6). As shown in Figure 5b, there are only 2.4% of PCGs (*n*=416) with isoforms belong to two or more categories, which is far less than the 19.3% of PCGs (*n*=3,917) with the similar type of isoforms in human (Uhlen et al. 2015). It is worth noting that the novel protein isoforms (86.74%) has a greater proportion of soluble proteins than known protein isoforms (71.49%). The results showed that the solubility of soluble proteins in liquids may be one of the reasons that due to some proteins were missed in current pig proteome.

**Figure 5.**
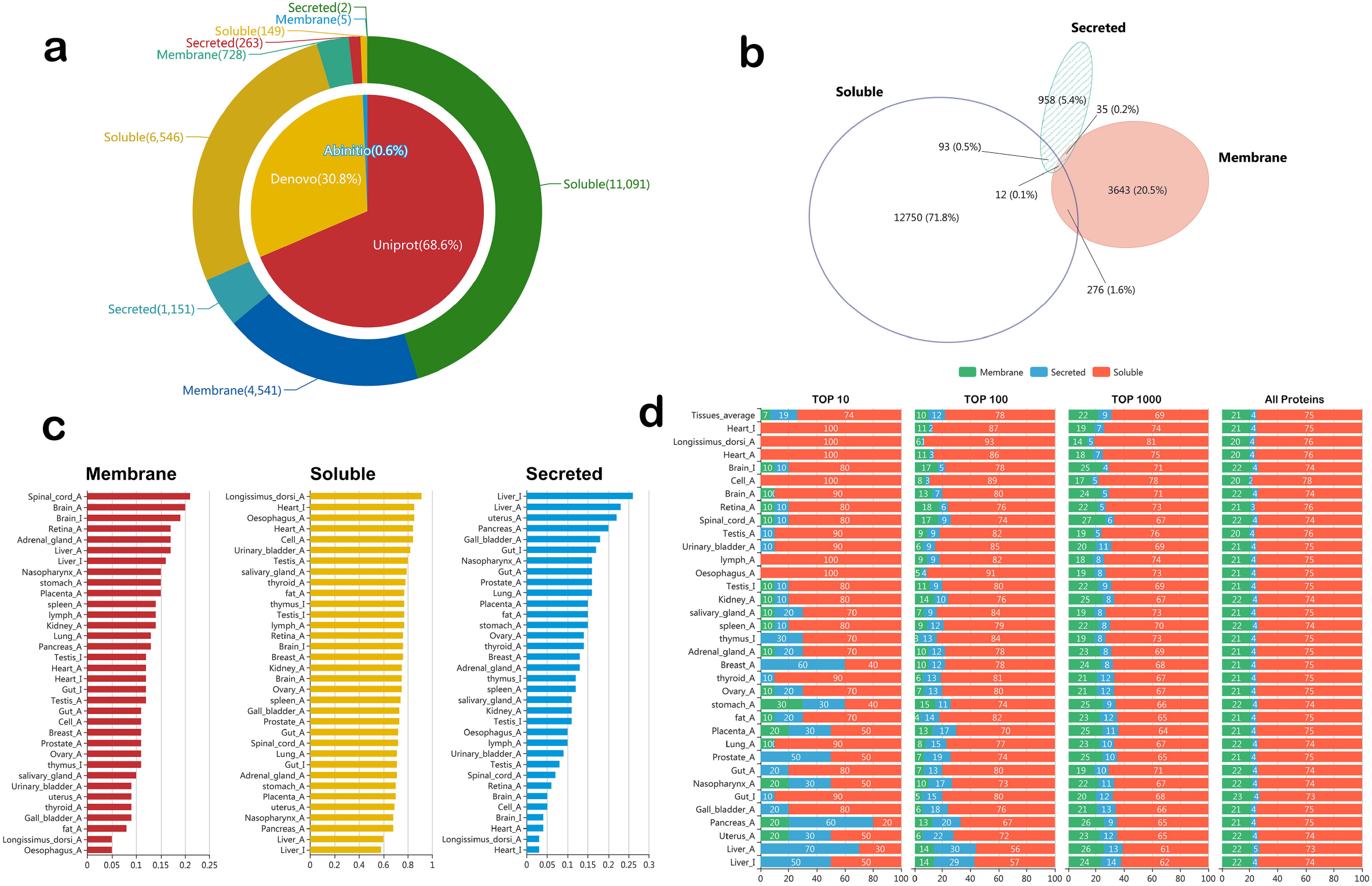
Classification of subcellular components within pig proteome. **a.** Pie charts for subcellular location of pig protein isoforms. Pie charts show the percentage of subcellular location for all pig protein isoforms. **b.** Venn diagram for subcellular location of pig proteins. Venn diagram reveals the number of genes in each of the three main subcellular location categories: membrane, secreted, and soluble. The overlap between the categories gives the number of genes with isoforms belonging to two or all three categories. **c.** The proportion of protein isoforms in 34 tissues to different subcellular components. **d.** The proportion of three subcellular components in 34 tissues. We respectively selected the levels of expression with top 10, top 100, top 1000 and all proteins as protein sets for each tissue.

More interestingly, we found that the organ or tissue function were also related to subcellular of their expressed proteins. Ranking all of identified proteins by their zFPKM value for each tissue, we selected the top 1% to represent their main proteins. As shown in Figure 5c, the higher proportion of membrane proteins were associated with nervous tissues, such as spinal cord, brain, retina. Besides, muscle tissues have a higher proportion in soluble protein, and the higher proportion of secreted proteins were represented by higher expression especially in some secretory tissues, such as liver, uterus, pancreas, gall bladder, gut *etc*. Besides, similar to human proteome(Uhlen et al. 2015), these highly expressed protein especially in secretory tissues were usually tend to the secreted components and representative to the tissue function (Figure 5d). For example, the *ALB* (albumin) gene codes a secreted protein with the highest expression seen in liver tissue of adult and infancy pig, with its main function being in the regulation of blood plasma colloid osmotic pressure. Whereby we further predicted the potential function of these unknown isoforms by referring to well-annotated gene that having the parallel expression pattern and subcellular components. For example, both of LOC100620249 and *PGC* (Progastricsin) genes were highly expressed secreted proteins in the stomach tissue, and the latter is a known secreted protein and constituting a major component of the gastric mucosa. This demonstrates the LOC100620249 more likely plays an important role in gastric mucosa of pig, which provide a valuable resource for enhancing studies of pig genomics and biology.

### Inferring orthologous functions of unknown pig proteome across multiple species

To pursue stronger evidence and orthologous functions for these unknown proteins, we further aligned the sequence of each isoform against the top 10 species databases. We adopted two criteria to identify homologous sequences to the newly identified swine proteins with those of other species: (1) percent identity is greater than 80%; (2) length of homologous sequence is longer than 80% of the swine protein sequence. Consequentially, 3,656 out of 7,319 (49.95%) unknown protein isoforms were inferred to have orthologous in other species. 90.29% orthologous isoforms (*n*=3,301) were identified in at least other two species, while 36.95% of orthologous isoforms (*n*=1,351) were the common isoforms for 9 (Chicken) or all 10 species (Figure 6a). Interestingly, Even the novel protein isoforms still have 41.72% of the orthologous protein isoforms (*n*=1,545) with other species, and almost 65.3% (*n*=1,009) of them were mapped in the pig genome (*Sus scrofa* 11.1) (Figure 6b). The result indicated that the exploited novel proteins herein can be considered as the reliable proteome data that significantly enhances both the pig genome annotation and the current pig protein database with convincing evidence.

**Figure 6.**
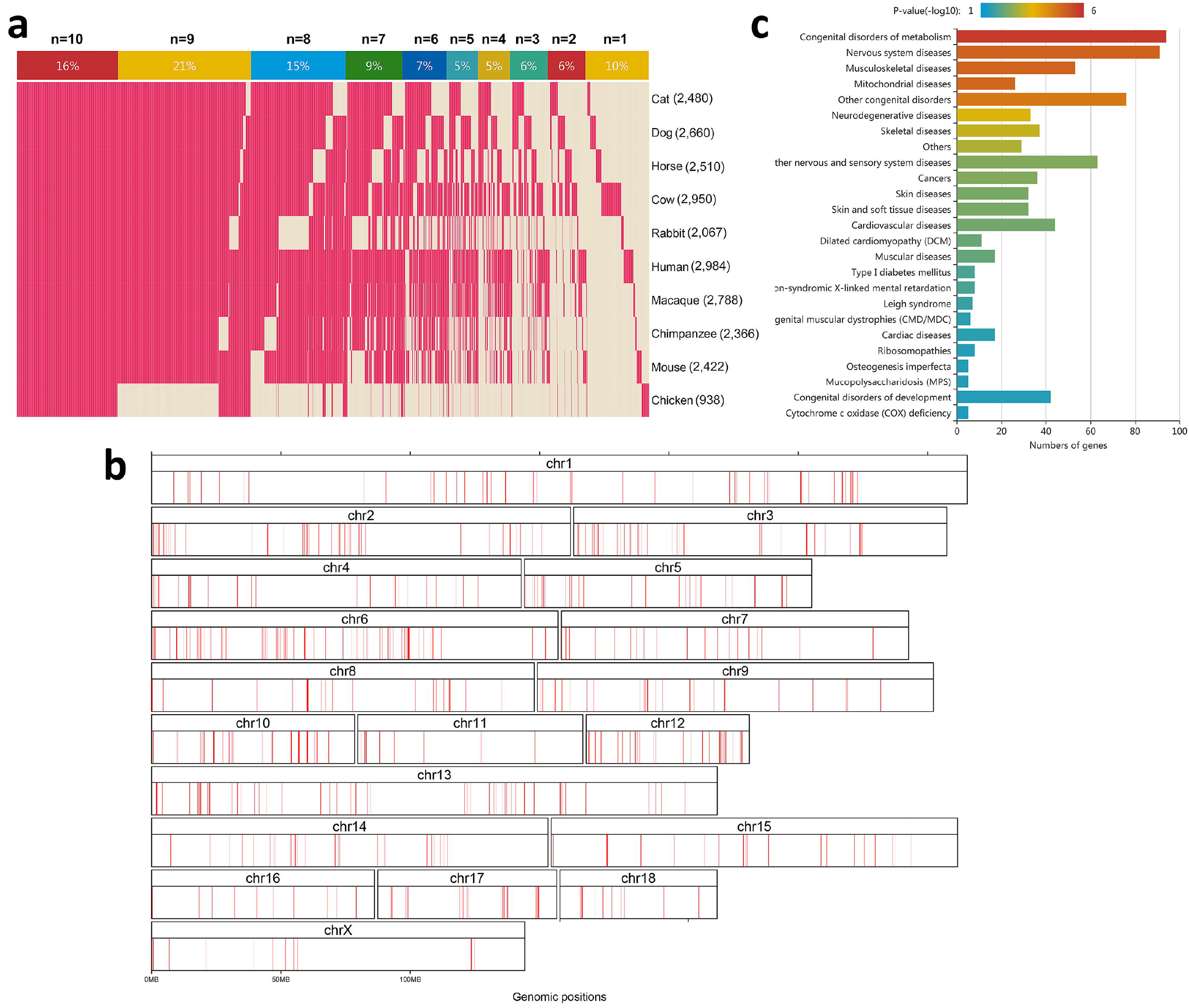
Orthologous of unknown pig proteome across multiple species. **a.** The heatmap for showing 3,656 orthologous isoforms among 10 species. For each isoform, the N represent the number of species that pigs shared homology with. The percentages within the colour bars mean the proportion of genes in all 3,656 homologous isoforms for each “N”. **b.** The distribution of the novel proteins in the pig genome. **c.** 25 KEGG disease pathway for human orthologous proteins of pig novel proteins.

In addition, compared with the existing orthologues in omabrowser (http://omabrowser.org) and current genome sequences, 3,656 of the unknown protein isoforms enriched 14,837 novel pairwise orthologous relationships between pigs and other species (Table S7). These pairwise orthologous relationships of proteins between pigs and other species provided a feasible way to investigate the potential function of corresponding PCGs in the pig genome if these homologous proteins have been well studied in other species. Therefore, considering the most complete set of annotated genes in human proteome, we preformed the functional gene set enrichment for human orthologous proteins of these unknown protein isoforms to speculate their potential function. A functional Gene Ontology (GO) analysis for these unknown protein isoforms showed that most of GO term describes of cell and intracellular part (corrected *P* < 0.01), which provide an important supplement to understand the biological process in pig. Meanwhile, by further examining the functional characterization of these unknown protein isoforms, we found 65 Kyoto Encyclopedia of Genes and Genomes (KEGG) pathways were represented in our unknown proteome, mainly involving metabolic pathways (corrected *P* = 7.2e^−19^), focal adhesion (corrected *P* = 3.1e^−9^), regulation of actin cytoskeleton (corrected *P* = 4.4e^−8^). Importantly, we found 25 disease signaling pathways from the KEGG disease database (corrected *P* < 0.05) that included the metabolism, nervous system, skeletal, muscular, skin diseases (Figure 6c). These findings will help us better recognize the potential function of the unknown pig protein isoforms, and provide a new insight into enhancing the value of pigs as a biomedical model for human medicine and as donors for porcine-to-human xenotransplantation.

## Discussion

Here we presented the landscape of a tissue-based proteome for pigs. Our findings not only offered the confirmatory evidence for 84.84% of existing pig proteins that have been deposited in the UniprotKB (*n* = 16,738), pig PeptideAtlas (*n* = 9,204) and NCBI Protein database (*n* = 17,781), but also identified 3,703 novel protein isoforms which missed in current pig proteome. Besides, we also detected 669 protein isoforms from uncharacterized LOC genes and 2,947 protein isoforms without NCBI annotation in the current reference *Sus scrofa1*1.1 genome. Eventually, a total of 7,319 unknown protein isoforms were exploited to further optimize the annotation of PCGs for the current pig genome.

We systematically characterized unknown protein proteome for their expression features, subcellular components and orthologous functions, providing a valuable resource for enhancing studies of pig genomics, as well as offering the opportunities for exploring the potential function of these unknown proteins. Our findings clearly showed that the missing protein annotation in previous studies was due to the two aspects: (1) low-quality assembly in *Sus scrofa*10.2 genome, and (2) the specific features that low expression levels, tissue specificity, and greater proportion of soluble components in novel protein isoforms. The in-depth identification and subcellular characterization of proteome using multiple tissues make it feasible to develop a tissue-based pig proteome map and facilitate studies of functional genomics and relevant fields. We effectively improved genome annotation for 5,027 unknown protein isoforms (excluding LOC genes) by mapping their protein sequence to current pig genome (*Sus scrofa*11.1), of which 4,819 were assigned to chromosomes and 208 were resided on 33 unplaced scaffolds.

High-resolution profiling of pig transcriptome allows us to further reveal 1,746 unknown protein isoforms that display a tissue- (1,694) or group-enriched (52) function expression pattern. Besides, 507 of unknown protein isoforms were ubiquitously expressed in 34 tissues, which raised 9.8% of the potential “housekeeping” gene in the pig genome. These findings provide new insight into understanding the molecular function of the respective tissue or organ. Further inferring the biological function of unknown pig proteome by human orthologous proteome, we found that these unknown protein isoforms were enriched in 65 KEGG pathways and 25 disease signaling pathways, including the pathways involved in disease of concern for human medicine, such as metabolism, nervous system, skeletal, muscular and skin diseases.

The integrated data of proteome and transcriptome in the 34 pig tissues herein were presented in Supplemental file 2, and 7,319 unknown protein isoforms with corresponding genomic locations, expression landscapes, subcellular characterizations, orthologous proteins and predicted functions were also summarized in Table S8. All findings herein will provide the valuable insight and resources for enhancing studies of pig genomics and biomedical model application to human medicine in the future.

## Conclusions

We have profiled a draft map of pig proteome and identified 7,319 unknown protein isoforms using 34 major normal pig tissues. Further we functionally annotated novel protein isoforms through profiling the pig transcriptome with high-throughput RNA sequencing (RNA-seq) of the same pig tissues, improving the genome annotation of corresponding protein coding genes. We predicted the tissue related subcellular components and potential function for these unknown proteins. Finally, we mined orthologous genes of unknown protein isoforms across multiple species and revealed important disease signaling pathways. Our study enhances the pig genome annotation and contributes to accelerating biomedical research for porcine-to human xenotransplantation.

## Methods and Materials

### Sample acquisitions

Pig tissue samples and PBMC used for protein identification and mRNA expression analyses were collected from the Ninghe breeding pig farm in Tianjin, China. For purpose of generating a profiles of transcriptome and proteome of all major organs and tissues in pig, we totally collected 34 samples (i.e., 33 pooled tissues and the PBMC) from the nine unrelated Duroc pigs, including three adult male pigs and three female pigs at 200 to 240 days of age, as well as three male piglets (infancy) at 21 to 25 days of age. All pig tissues were histologically confirmed to be normal and healthy by an experienced pathologist. An overview of all involved tissue and cell samples is provided in Table S1.

### Preparation pig samples

All samples were snap frozen within the first 20 minutes after slaughter and stored in liquid nitrogen (−196°C) until usage. PBMC were isolated using Ficoll-Hypaque PLUS (GE Healthcare), following the manufacturer’s instructions. In brief, the whole blood was first diluted by an equal volume of phosphate buffer solution (PBS). Then, 20 ml of diluted blood was carefully added on top of 10 ml of Ficoll-Hypaque solution in a 50 ml conical tube and centrifuged at 460 g for 20 min at room temperature. After centrifugation, the middle whitish interface containing mononuclear cells was transferred to a new tube, and washed by PBS followed by centrifugation at 1000 rpm for 10 min twice.

### Separation of protein and RNA

Fresh frozen tissue was thawed, cut into small pieces and extensively washed with precooled phosphate buffered saline. A pool of equal amounts of tissues from three unrelated pigs was homogenized and sonicated in cold lysis buffer. Extraction of 100 μg protein using protein extraction buffer (8 M urea, 0.1% SDS) containing an additional 1 mM phenylmethylsulfonyl fluoride (Beyotime Biotechnology, China) and protease inhibitor cocktail (Roche, USA) was kept on ice for 30 min and then centrifuged at 16,000 × g for 15 min at 4 °C. The supernatant was collected and determined with BCA assay (Pierce, USA) and 10-20% SDS-PAGE. The cell lysate was stored at −80°C before LC-MS analysis.

Total RNA was purified from pooled tissues via the Trizol method (Invitrogen, Carlsbad, CA) according to standard protocols. RNA degradation and contamination was monitored on 1% agarose gels. The purity and contamination of total RNA was checked using NanoPhotometer^®^ trophotometer (IMPLEN, CA, USA) and Qubit^®^ RNAAssay Kit in Qubit^®^ 2.0 Flurometer (Life Technologies, CA, USA). RNA integrity was measured using the RNA Nano 6000 Assay Kit of the Bioanalyzer 2100 system (Agilent Technologies, CA, USA). All pig samples that met the criteria of having an RNA Integrity Number (RIN) value of 7.0 or higher and at least 5 μg of total RNA, were included and batched for RNA sequencing.

### Library construction and RNA sequencing

Total RNA of samples meeting quality control (QC) criteria were rRNA depleted and depleted QC was done using the RiboMinus™ Eukaryote System v2 and RNA 6000 Pico chip according to the manufacturer’s protocol. RNA sequencing libraries were constructed using the NEBNext^®^ Ultra™ RNA Library Prep Kit (Illumina) with 3μg rRNA depleted RNA according to the manufacturer’s recommendation. RNA-seq library preparations were clustered on a cBot Cluster Generation System using HiSeq PE Cluster Kit v4 cBot (Illumina) and sequenced using the Illumina Hiseq 2500 platform according to the manufacturer’s instructions, to a minimum of 10G reads per sample (corresponding to 125 bp paired-end reads). The sequenced RNA-Seq raw data for the 34 pig tissues is available from NCBI Sequences Read Archive with the BioProject number: PRJNA392949.

### Fractionation of peptide mixture using a C18 column

Peptide mixture from each sample was first lyophilized and reconstituted in buffer A (2% ACN, 98% H2O, pH10). Then, it was loaded onto a Xbridge PST C18 Column, 130 Å, 5 μm, 250 × 4.6 mm column (Waters, USA), on the DIONEX Ultimate 3000 HPLC equipped with a UV detector. Mobile phase consists of buffer A and buffer B (90% ACN, 10% H2O, pH10). The column was equilibrated with 100% buffer A for 25 minutes before sample injection. The mobile phase gradient was set as follows at a flow rate of 1.0 mL/minute: (a) 0−19.9 min: 0% buffer B; (b) 19.9–20 min: 0–4% buffer B; (c) 20–22 min: 4–8% buffer B; (d) 22–42 min: 8−20% buffer B; (e) 42–59 min: 20–35% buffer B; (f): 59–60 min: 35–45% buffer B; (g): 60–61min: 45–95% buffer B; (h) 61–66 min: 95% buffer B; (i) 66–67 min: 95–0% buffer B; (j) 67–91: 0% buffer B. A fraction was collected every minute from 24 min to 63 min, and a total of 40 fractions collected were then concentrated to 20 fractions, vacuum dried and stored at −80°C until further LC-MS/MS analysis.

### Liquid chromatography tandem mass spectrometry

Peptide mixture was analyzed on a Q Exactive instrument (Thermo Scientific, USA) coupled to a reversed phase chromatography on a DIONEX nano-UPLC system using an Acclaim C18 PepMap100 nano-Trap column (75 μm × 2 cm, 2 μm particle size, Thermo Scientific, USA) connected to an Acclaim PepMap RSLC C18 analytical column (75 μm × 25 cm, 2 μm particle size, Thermo Scientific, USA). Before loading, the sample was dissolved in sample buffer, containing 4% acetonitrile and 0.1% formic acid. Samples were washed with 97% mobile phase A (99.9% H2O, 0.1% formic acid) for concentration and desalting. Subsequently, peptides were eluted over 85 min using a linear gradient of 3-80% mobile phase B (99.9% acetonitrile, 0.1% formic acid) at a flow rate of 300 nL/min using the following gradient: 3% B for 5 min; 3–5% B for 1 min; 5-18% B for 42 min; 18–25% B for 11 min; 25–30% B for 3 min; 30–80% B for 1 min; hold at 80% B for 5 min; 80–3% B for 0.5 min; and then hold at 3% B for 21.5 min. High mass resolution and higher-energy collisional dissociation (HCD) was employed for peptide identification. The nano-LC was coupled online with the Q Exactive mass spectrometer using a stainless steel emitter coupled to a nanospray ion source. The eluent was sprayed via stainless steel emitters at a spray voltage of 2.3 kV and a heated capillary temperature of 320°C. The Orbitrap Elite instrument was operated in data-dependent mode, automatically switching between MS and MS2. Mass spectrometry analysis was performed in a data dependent manner with full scans (350-1,600 m/z) acquired using an Orbitrap mass analyzer at a mass resolution of 70,000 at 400 m/z on Q Exactive using an automatic gain control (AGC) target value of 1×106 charges. All the tandem mass spectra were produced by HCD. Twenty most intense precursor ions from a survey scan were selected for MS/MS from each duty cycle and detected at a mass resolution of 17,500 at m/z of 400 in Orbitrap analyser using an AGC target value of 2×105 charges. The maximum injection time for MS2 was 100 ms and dynamic exclusion was set to 20s.

### Validation of identified Proteins

In total, 71 peptides from 31 proteins (7 known proteins, 11 homologous novel proteins, 13 non-homologous novel proteins) were randomly selected for peptide synthesis (GL biochem) for validation of identified proteins. The synthesized peptide sequences were mixed and were processed twice by chromatographic separation using the Thermo EASY-nLC HPLC system and Thermo scientific EASY column. Mass spectral analysis was then performed by Q-Exactive (Thermo Scientific) and processed by Mascot V2.2. Finally, all these peptides were compared with those identified from our proteome analysis to verify novel proteins.

### QC processing

We conducted a quality control step on raw fastq reads for efficient and accurate RNA-seq alignment and analysis. In this step, raw reads were cleaned up for downstream analyses using the following steps: BBDuk (http://sourceforge.net/projects/bbtools/)(Bushnell 2014) automatically detected and removed adapter sequences; FASTQC (http://www.bioinformatics.babraham.ac.uk/projects/fastqc/) calculated the Q20, Q30 and GC content of the clean data for quality control and filtering; FASTX-Toolkit (http://hannonlab.cshl.edu/fastx_toolkit/) carried out homopolymer trimming to the 3’ end of fragments and removed the N bases from the 3’ end.

### Read mapping and assembly

RNA-seq data were mapped and genome indexed with Hisat 0.1.6-beta 64-bit(Kim et al. 2015) to t he pig genome release version *Sus scrofa11.1* (ftp://ftp.ncbi.nlm.nih.gov/genomes/all/GCF/000/003/025/GCF_000003025.6_Sscrofa11.1/). *Sus scrofa11.1* annotation was used as the transcript model reference for the alignment, splice junction identification and for all protein-coding gene and isoform expres sion-level quantifications. To obtain expression levels for all pig genes and transcripts across all 34 samples, FPKM values were calculated using Stringtie 1.0.4 (Linux_x86_64)(Pertea et al. 2015). A gene or transcript was defined as expressed if it’s FPKM value was measured less than 0.1 across all tissues. For each tissue, we applied zFPKM normalization method(Hart et al. 2013) to generate high-confidence estimates of gene expression.

### zFPKM level-based classification of genes

Refer to the gene classification in human, we also classified the pig genes into one of three categories based on the zFPKM levels in 34 samples: (1). “Tissue enriched” – only detected in single tissue, as well as at least 5-fold higher at the zFPKM level in one tissue compared to the tissue with a second highest expression. (2). “Group enriched” – the gene detected in all tissues from a groups, and the expression of genes in any tissue from the groups is higher than the tissue that not from the group. (3). “Expressed in all tissues” – detected in all 34 tissues.

### Construction of a reference protein database

To identify novel protein and improve existing proteins annotations in the pig genome, the database for protein searching (MS/MS data searched against protein database) was taken from four different levels using in-house perl scripts, including: (1). UniProt database (Sus scrofa) (2). Three-frame-translated mRNA de novo sequences from the current study (3). Six-frame-translated pig genome database. The details are as follows: Primary database of proteins: Resource protein data sets for pig (UniProt version 20150717 containing 34,131 entries, with 1,486 Swiss-Prot, 32,643 TrEMBL) were downloaded from the UniProt database (http://www.uniprot.org/).

Secondary database of proteins: It is well known that pig proteins of insufficiently represented by detectable known proteins, because of the incomplete nature of the pig genome assembly and limited annotation. In our study, three RNA resources were used (Table S9): (1). EST datasets including 34,131 entries from UCSC (http://hgdownload.soe.ucsc.edu/goldenPath/susScr3/bigZips/) and 1,676,406 entries from the NCBI database (http://www.ncbi.nlm.nih.gov/nucest). ESTs are normally assembled into longer consensus sequences for three-frame-translated mRNA protein database using iAssembler version 1.3.2.x64(Zheng et al. 2011) with default parameter. (2). Paired-End (PE) read libraries including 34 RNA sequencing libraries from our study and 7 previously published article and NCBI database. To construct a complete protein database for three-frame-translated mRNA, we used Trinity (version 2.0.6)(Grabherr et al. 2011) for *de novo* transcriptome assembly from RNA-Seq data, and identified potential coding regions within Trinity-reconstructed transcripts by TransDecoder (developed and include with Trinity). (3). Single-end (SE) reads from 10 previous studies were downloaded from NCBI (http://www.ncbi.nlm.nih.gov/sra/). The method for sequence assembly and coding region prediction were similar to that used for the paired-end (PE) reads.

Tertiary database of proteins: To capture the proteins missed during the laboratory discovery process as far as possible, protein annotation of the pig genome was carried out using the ab initio methods with GeneScan software(Ramakrishna and Srinivasan 1999). Finally, we used BLASTP to identify proteins and remove duplicates between different protein databases (Retention order: UniProt > De novo > Ab initio).

### Peptide identification based on database searching

All MS/MS samples were analyzed using Mascot (Matrix Science, London, UK; version 2.5.1)(Sadeh et al. 1999) and X!Tandem (The GPM, thegpm.org; version CYCLONE (2010.12.01.1))(Craig and Beavis 2004). Mascot was set up to search the pig databases (UniProt, de novo, Assembly, ab initio database) and the cRAP database (common Repository of Adventitious Proteins; download date: 07 Jul 2015; 116 sequences) assuming the digestion enzyme trypsin.

The high resolution peaklist files were converted into Mascot generic format (mgf) prior to database searching. X!Tandem was set up to search a subset of the pig databases, also assuming trypsin. The target-decoy option of Mascot and X!Tandem were enabled (decoy database with reversed protein sequences). Mascot and X!Tandem were used to search with a fragment ion mass tolerance of 0.050 Da and a parent ion tolerance of 10.0 PPM. Carbamidomethyl of cysteine was specified in Mascot and X!Tandem as a fixed modification. Gln- > pyro-Glu of the n-terminus, oxidation of methionine and acetyl of the n-terminus were specified in Mascot as variable modifications. Glu- > pyro-Glu of the n-terminus, ammonia-loss of the n-terminus, Gln- > pyro-Glu of the n-terminus, oxidation of methionine and acetylation of the n-terminus were specified in X! Tandem as variable modifications.

Scaffold (version Scaffold_4.4.5, Proteome Software Inc., Portland, OR) was used to validate MS/MS based peptide and protein identifications. Peptide identifications were accepted if they achieved an FDR < 1% by the Scaffold Local FDR algorithm. Protein identifications were accepted if they had an FDR < 1% and contained at least 2 identified peptides. Protein probabilities were assigned by the Protein Prophet algorithm. Proteins that contained similar peptides and could not be differentiated based on MS/MS analysis alone were grouped to satisfy the principles of parsimony. Proteins sharing significant peptide evidence were grouped into clusters. In the database searching workflow, unmatched MS/MS spectra generated from the Uniprot database search were then searched against next level protein database (De novo, Ab initio).

### Mapping the protein isoforms to the pig genome

We attempted to map all unknown protein isoforms against the pig genome using MAKER annotation workflow(Cantarel et al. 2008). First, the low-complexity repeats of pig reference genome were soft-masked by RepeatMasker. Then, the unknown protein isoforms (without LOC genes) were aligned to the masked reference genome by BLAST(Mount 2007) for identifying their genomic location roughly. Last, Exonerate(Slater and Birney 2005) was used to realign and polish the exon–intron boundaries of the unknown gene with the splice-site aware alignment algorithm.The house-python script being used to deal with the result: if a successfully aligned protein had 95% identity overall, 95% coverage and the distance from its neighboring exon being less than 50Kb, it was recorded to be an effectively aligned sequence.

### Subcellular prediction and classification of pig proteome

The prediction of pig membrane proteins was carried out similarly to how these proteins were classified in the human proteome. A total of seven methods were used to identify membrane protein topology with different assessment algorithms, for example, topological models, neural networks, support vector machines (SVMs), scale of free energy contributions and hidden Markov models (HMMs): MEMSAT3(Jones 2007), MEMSAT-SVM(Nugent and Jones 2009), SPOCTOPUS(Viklund et al. 2008), THUMBUP(Zhou and Zhou 2003), SCAMPI multi-sequence-version(Bernsel et al. 2008), TMHMM(Sonnhammer et al. 1998), and Phobius version 1.01(Kall et al. 2004). In our study, the proteins were assigned as transmembrane if they were predicted by at least four out of the seven methods.

In accordance with human secretome analysis, the prediction of signal peptides (SP) was based on Neural Networks and Hidden Markov models with three software programs: SignalP4.0(Petersen et al. 2011), SPOCTOPUS and Phobius version 1.01. The proteins, predicted by at least two out of the three methods, to contain a signal peptide were classified as potentially secreted.

Integrating the prediction of pig membrane proteins and prediction of pig secretome proteins, we classified each pig protein into one of three classes: secreted, membrane or soluble (neither membrane nor secreted protein). In order to compare the proteome between pig and human conveniently, we also constructed four major categories for classifications of the protein-coding genes with multiple protein isoforms: (1). “soluble” just containing soluble category (2). “secreted” were combined with the soluble/secreted and the secreted (3). “membrane” including soluble/membrane and the membrane groups (4). “membrane and secreted isoforms” containing secreted/membrane and soluble/secreted/membrane groups.

### Weighted gene co-expression network analysis

In order to reveal the groups of protein coding genes that are functionally related in the whole pig organism, 34 pig tissue data sets were constructed using the WGCNA method. In our study, we mainly used the blockwiseModules function in the WGCNA R package(Langfelder and Horvath 2008) to perform the coexpression network construction, with the following parameters: corType = pearson, maxBlockSize = 30,000, power = 8, minModuleSize = 30, mergeCutHeight = 0.1. The brief function of blockwiseModules automatically constructed a correlation network, created a cluster tree, defined modules as branches, merged close modules, and yielded the module colors and module eigen genes for subsequent analysis (such as visualization by the plotDendroAndColors function).

### Functional annotations for pig PCGs

Gene ontology (GO) analysis and KEGG (http://www.genome.jp/kegg/) pathway enrichment analysis were performed with KOBAS 3.0 (http://kobas.cbi.pku.edu.cn/anno_iden.php). GO terms appearing in this study are summarized within three categories: cell component, molecular function and biological process. In view of the most complete genes annotation in human genome, we gave priority to those human annotated genes which were homologous to pig genes and utilized them as the background.

## Availability of data and material

The sequenced RNA-Seq raw data for the 34 pig tissues is available from NCBI Sequences Read Archive with the BioProject number: PRJNA392949. The pig proteomic data was deposited in PRIDE with the Project Accession was PXD006991.

## Additional files

**Figure S1-14: Quality assessment of protein identification.**

S1-S3. Density distribution for number of unique peptide at the region from 0 to 10, 0 to 20 and 0 to 50 respectively.

S4-S6. Density distribution for number of unique spectrum at the region from 0 to 10, 0 to 20 and 0 to 50 respectively.

S7-S9. Density distribution for number of spectrum counts at the region from 0 to 10, 0 to 20 and 0 to 50 respectively.

S10. Density distribution for identification probability.

S11. The bar plot for number of protein with different peptide bins.

S12. The bar plot for number of protein with different coverage bins among 34 tissues.

S13. The bar plot for number of protein with different tissues among 10 coverage bins.

S14. The bar plot for number of protein with different coverage bins.

**Supplemental tables: Table S1-S9 (XLSX)**

Table S1 Validation of 71 peptides from 31 proteins

Table S2 Overview of alignment within 34 tissues Table S3 Gene expression patterns in 34 tissues

Table S4 The co-expression interactions of 6,163 ubiquitously expressed gene Table S5 Subcellular location of the pig proteome (isoform)

Table S6 Subcellular location of the pig proteome (protein)

Table S7 Details of homologous protein with 10 species

Table S8 Overview of functionally predictive resource for 7,319 unknown protein isoforms Table S9 RNA-seq resource tables

**Supplemental file 1**: Validation of identified proteins: MS/MS spectra from 71 synthetic peptides with those obtained from analysis of pig tissues.

**Supplemental file 2**: The fasta file of 3,703 novel protein isoforms.

## List of Abbreviations

RNA-seq: : RNA sequencing
PCGs: : protein-coding genes
LC-MS/MS: : liquid chromatography tandem mass spectrometry
EST: : expressed sequence tag
PSMs: : peptide spectrum matches
FDR: : false discovery rate
PBMC: : peripheral blood mononuclear cells
GO: : Gene Ontology
KEGG: : Kyoto Encyclopedia of Genes and Genomes
RIN: : RNA Integrity Number

## Ethics approval and consent to participate

All procedures involving animals were performed by a licence holder in accordance with the protocol approved by the Institutional Animal Care and Use Committee (IACUC) of China Agricultural University (Beijing, People’s Republic of China, permit number: DK1023).

## Consent for publication

Not applicable.

## Authors’ contributions

J-F.L. conceived and designed the experiments. P.Z. performed transcriptome and proteome analyses. C.D., H.K. and L.Z. performed pathway analysis and graphic design. X.Z., J.L., C.N., H.W. and Y.C. collected samples and prepared for sequencing. X.Z. and W.F. assisted the experimental validations. J-F.L., P.Z., J.S., Z.H., G.L., W.W., Y. Y., B.L., and J.S. wrote and revised the paper. All authors read and approved the final manuscript.

## Acknowledgements

We thank Ziyao Fan, Yichun Dong and Kaijie Yang for samples collection.

